# Optimal Multichannel Artifact Prediction and Removal for Brain Machine Interfaces and Neural Prosthetics

**DOI:** 10.1101/809640

**Authors:** Mina Sadeghi Najafabadi, Longtu Chen, Kelsey Dutta, Ashley Norris, Bin Feng, Jan WH Schnupp, Nicole Rosskothen-Kuhl, Heather Read, Monty Escabi

**Affiliations:** Department of Electrical and Computer Engineering, University of Connecticut, Storrs, CT 06269; Department of Biomedical Engineering, University of Connecticut, Storrs, CT 06269; Department of Psychology, University of Connecticut, Storrs, CT 06269; Department of Biomedical Sciences, City University of Hong Kong, Hong Kong SAR; Neurobiological Research Laboratory, Section for Clinical and Experimental Otology, University Medical Center Freiburg, Freiburg, Germany

**Keywords:** Artifact removal, electrical stimulation, nerve fibers, cochlear implants, neural implant, brain machine interface, Wiener filter

## Abstract

Neural implants that electrically stimulate neural tissue such as deep brain stimulators, cochlear implants (CI), and vagal nerve stimulators are becoming the routine treatment options for various diseases. Optimizing electrical stimulation paradigms requires closed-loop stimulation using simultaneous recordings of evoked neural activity in real time. Stimulus-evoked artifacts at the recording site are generally orders of magnitude larger than the neural signals, which challenge the interpretation of evoked neural activity. We developed a generalized artifact removal algorithm that can be applied in a variety of neural recording modalities. The procedure leverages known electrical stimulation currents to derive optimal filters that are used to predict and remove artifacts. We validated the procedure using paired recordings and electrical stimulation from sciatic nerve axons, high-rate bilateral CI stimulation, and concurrent multichannel stimulation in auditory midbrain and recordings in auditory cortex. We demonstrate a vast enhancement in the quality of recording even for high-throughput multi-site stimulation with typical improvements in the signal-to-noise ratio between 20-40 dB. The algorithm is efficient, can be scaled to arbitrary number of sites, and is applicable in range of recording modalities. It has numerous benefits over existing approaches and thus should be valuable for emerging neural recording and stimulation technologies.

## INTRODUCTION

Brain machine interfaces and neural prosthetics, such as cochlear implants (CIs) and vagal nerve stimulation, increasingly rely on stimulation of neural tissue and concurrent neural recordings to either assess neural function, optimize electrical stimulation, or to provide neural feedback [1-4]. In such applications, capacitive and inductive coupling of the delivered electrical currents within the neural tissue produces artifacts at distant recording sites that are often orders of magnitude larger than the recorded neural activity. Such artifacts obscure neural activity and make it difficult to interpretation and quantification of neural data. Such artifacts are also widely present in multi-channel electrophysiological recordings that use electrical stimulation to characterize neural function and in clinically relevant signals that are measured during concurrent stimulation, such as EEG and ECoG. Artifact removal from neural recordings is thus necessary to isolate neural responses to the electrical stimuli in order to asses neural encoding, neural transformations, for clinical assessment, and to optimizing the stimulation efficiency of brain machine interfaces and prosthetic devices.

Existing techniques for artifact removal invariably focus on the recorded artifact waveforms using artifact removal algorithms that are *blind* to the artifact generator sources. That is, nearly all approaches assume that the electrical stimulation currents are unknown, which in most instances is not the case since they are being delivered by the experimenter via a computer interface. These include artifact template subtraction [5-8], local curve fitting [9], sample-and-interpolate technique [10], and independent component analysis [11-16]. Such waveform centric algorithms can suppress artifacts in certain stimulation/recording paradigms, yet they often rely on artifact estimation and subtraction techniques that place assumptions on the statistics of the neural signals and artifacts and can themselves distort the target neural signals. A drawback of many such techniques is that they typically require that the recorded artifacts arise from a single isolated source so that the artifacts are reproducible and non-overlapping in time, which is often not the case. Furthermore, such techniques are often difficult to implement in real-time with realistic current stimulation paradigms and are instead used for post-hoc removal of artifacts. Most current methods also fail with multi-channel stimulation (i.e., multiple sources), however, a recent method used Gaussian mixture models to improve the quality of spike sorting during multi-site stimulation and concurrent multi-site recordings [17]. As for other techniques, such a method places statistical constraints on the nature of both the neural activity and artifact signals and is not directly applicable to the analysis of continuous neural signals, such as ECoG and EEG. Furthermore, the technique does not make use of known electrical stimulation currents which can substantially enhance the artifact removal as we will demonstrate. Finally, many current methods fail when multiple artifacts are generated in close succession during fast current stimulation, such as for CIs where hundreds to thousands of pulses per second, often overlapping in time, are delivered across multiple stimulation electrodes [18]. In such cases, artifact removal can be improved by decreasing the rate of stimulation, although this leads to abnormal stimulation regimes not compatible with real time neural feedback or for characterizing natural neural processing with such devices [18].

Here we develop an optimal multichannel artifact removal algorithm that can be applied during high-throughput multi-site electrical stimulation. Unlike nearly all artifact removal procedures which are *blind* and assume that electrical currents are unknown, the procedure capitalizes on the fact that transformation between electrical stimulation and recording arrays arises through *linear* capacitive and inductive coupling [19] and the fact that stimulation currents are actually known *a priori* in most instances. This allows us to derive optimal linear filters to model the transformation between each of the stimulation and recording electrode pairs. The optimal filters are then used to predict the neural recording artifacts that arise from individual input channels, which can then be removed via subtraction. The procedure is versatile and can be applied to a variety of neural recording modalities including single, multi-unit, and continuous field potential recordings. Furthermore, because the algorithm estimates the transfer functions between every stimulation and neural recording electrodes the procedure can be applied irrespective of the stimulation currents used and is thus compatible with single and multi-site stimulation, high-rate stimulation, and is applicable to electrical stimuli with arbitrary pulse amplitudes and shapes. By applying the procedure to sample neural datasets (single and multi-channel stimulation), we demonstrate a vast signal-to-noise ratio improvement ranging between 20-40 dB.

## MATERIALS AND METHODS

### Multi-Input Multi-Output Artifact Prediction Wiener Filter

We develop an optimal Wiener filter algorithm to predict neural recording artifacts upon delivering electrical stimulation currents on a multi-channel stimulating electrode array. This predicted artifact is then subtracted from the actual neural recording trace to yield a noise free (or noise reduced) estimate of the neural activity.

We assume a generalized multi-input (stimulation) multi-output (recording) framework for developing a linear filter approximation of the recording artifact. This framework is also applicable and extends to the single input-single output case (single recording and stimulation channels). Given that electrical stimulation artifacts are the result of linear capacitive and inductive coupling between the neural tissue and the stimulating and recording electrodes [19], we model the transformations between the electrical stimulus and recorded artifact as a linear filter with unknown impulse response (or equivalently transfer function). Each stimulating and neural recording electrode pair has its own electrical characteristics and thus a unique impulse response to be determined based on the input and output data. Also, given that such electrical coupling is linear, the composite multi-site stimulation artifact is modeled as a linear sum of the artifacts generated by each stimulation channel and thus we have:

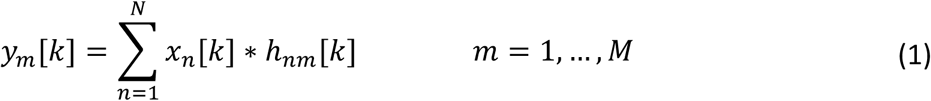

in which *y*_*m*_ [*k*] is the predicted artifact for channel *m* (**y**_*m*_ in vector form), *h*_*nm*_ [*k*] is the impulse response between the n-th stimulation channel and m-th neural recording channel (**h**_*nm*_ in vector form), and *x*_*n*_ [*k*] is the electrical stimulation signal applied to stimulation channel *n* (**x**_*n*_). In matrix form **y** = **hx** where **y** = [**y**_1_ … **y**_*M*_] is a matrix containing the predicted outputs for the *M* recording channels, **x** = [**x**_1_ … **x**_*N*_] is a matrix containing the input electrical stimulation signals across *N* stimulation channels, and

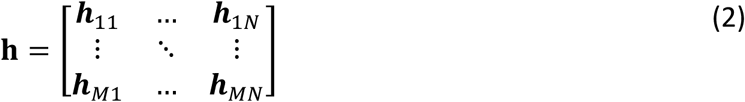

is a matrix containing the impulse responses vectors (***h***_nm_) between all stimulation and recording channels.

The goal is to derive the filter matrix **h** using experimental measurements. The estimated filter can then be used to predict the recorded artifacts. The optimal solution is obtained via the Wiener-Hopf equation [20]

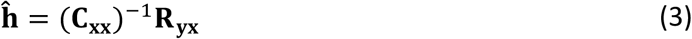

where **ĥ** is the filter matrix solution that minimizes the mean squared error between the predicted and real artifacts,

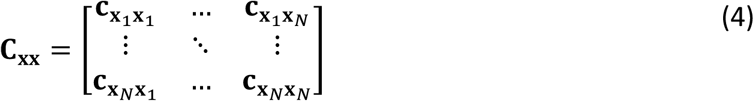

represents the stimulation signal covariance matrix which contains correlation functions 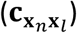 between the *n*-th and *l*-th (*l, n* = 1, …, *N*) input channels, and

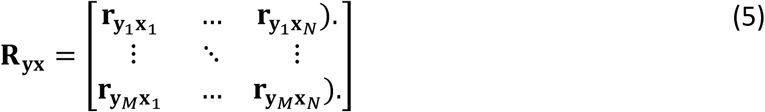

is a matrix containing the cross-correlation functions between the *m*-th output and *n*-th input channels 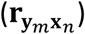.

Upon deriving the multi-site filters, the stimuli artifacts are then predicted by convolving the measured filters with the given input signals as in Eqn. 1. Finally, the predicted artifacts are subtracted from the recorded data yielding the noise reduced estimate of the neural traces. Although Eqn. 3 is derived for Multi-Input Multi-Output (*N* > 1, *M* > 1) neural recording and stimulation scenarios in mind, the procedure generalizes and is also compatible with Multi-Input Single-Output (*N* > 1, *M* = 1), Single-Input Multi-Output (*N* = 1, *M* > 1), and Single-Input Single-Output (*N* = 1, *M* = 1) neural stimulation and recording.

As a note, we point out that the form of the *predictive* Wiener filter used here differs from *blind deconvolutional* Wiener filters used previously for artifact removal which assume that the artifact generating signals are unknown [21-23]. Deconvolutional filters use the signal and noise spectrum statistics to optimally reject the artifact signal via deconvolution. In general, because the signal and noise spectrums often overlap such approaches tend to distort the neural signals of interest upon removing the artifacts and are not intended to fully remove the artifact. In our case, the wiener filter is instead used to predict the recorded artifact from *known* inputs, which can then be removed from the neural recording by subtraction without distorting the neural signal.

### Signal to Noise Ratio Estimation

We used a shuffled trial procedure to *estimate* the artifact (noise) and neural signal power spectrums which we then use to estimate the signal-to-noise ratio (SNR) of the neural recording or the artifact reduction ratio (ARR). The procedure requires that we deliver an identical electrical stimulation signal from two trials in order to estimate the signal and noise power spectrum. Consider a recorded neural trace

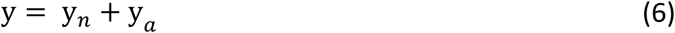

where y_*n*_ represents the noiseless neural trace (i.e., no artifact) and y_*a*_ represents the recorded artifact. If y’ represents the data recorded in the second trial of a repeated experiment (i.e., same electrical stimulation signal) then the artifact should be identical between the two trails (y_*a*_) so that

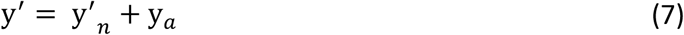

where here, y’_*n*_, is the neural response component for the second trial. This component differs from the first trial response (y_*n*_) because of neural variability. Computing the cross-spectral density (CSD) between the two trails yields

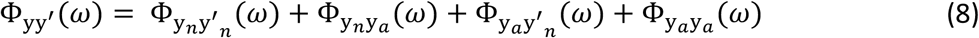

Similarly, the power spectral density (PSD) of the first trial is

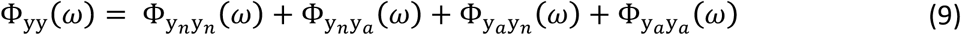

Given that the artifact signal is reproducible across trials and typically much larger than the recorded neural activity (e.g., as seen for the examples of Figs. 1-4), the artifact term in Eqn. 8 dominates

**Figure 1:**
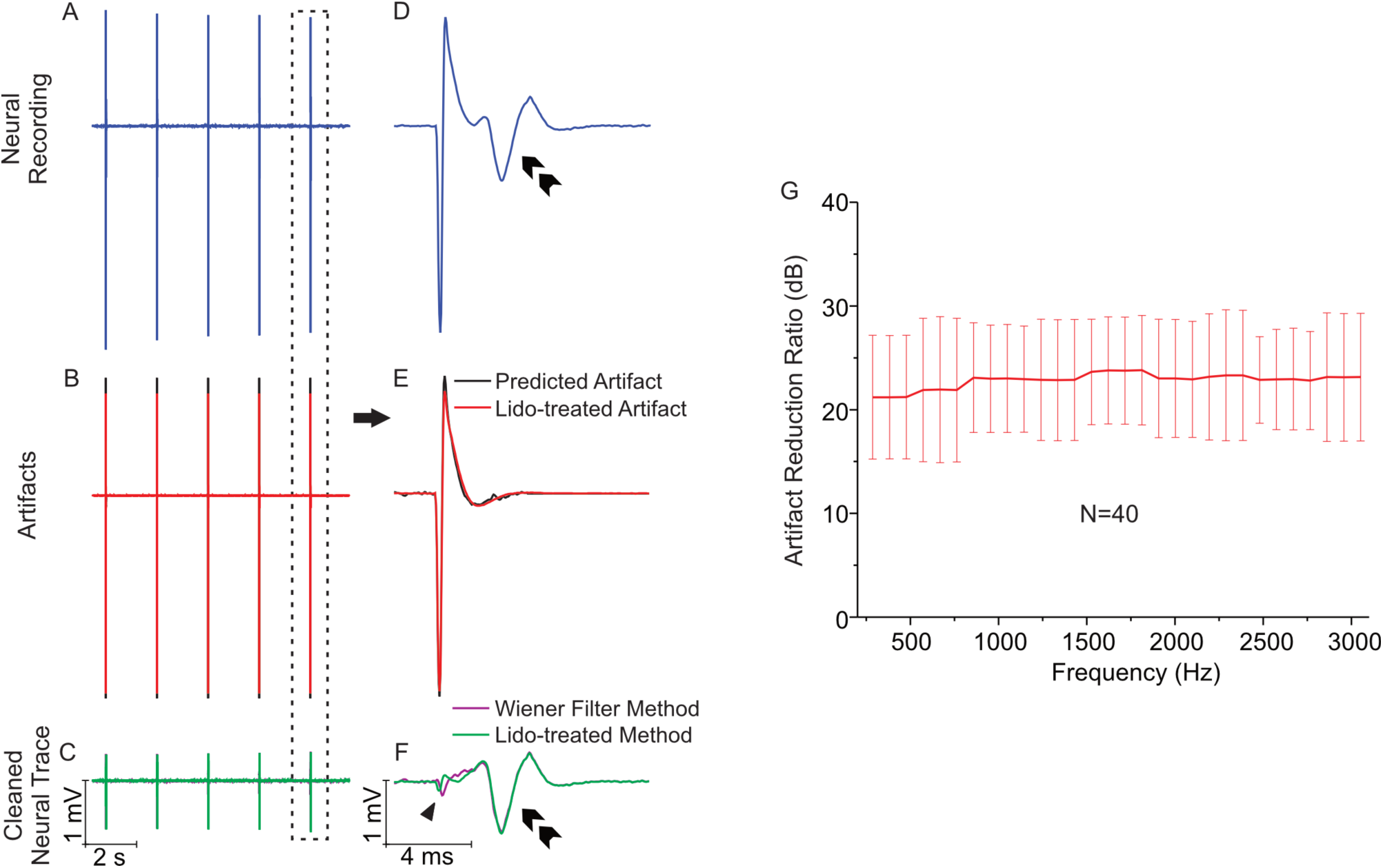
Artifact removal from neural recordings in mouse sciatic nerve. (A) shows recorded neural signals evoked by electrical stimulations (−320 uA peak amplitude, 0.2 ms duration); (B) illustrates lidocaine-treated artifacts (red) and filter predicted artifacts (black) of the corresponding stimulus; (C) is the cleaned neural activity (A) after the subtraction of lidocaine-treated artifacts (green) and predicted artifacts (purple) from neural recordings. (D), (E) and (F) are the magnified views of a single stimulation event in (A), (B) and (C), respectively. (G) shows the artifact reduction ratio. The amplitude of stimulus artifacts was significantly reduced after subtraction by the Wiener filter predicated artifacts, achieving an average artifact reduction ratio of 22.8 dB across the frequency range (0.3 to 3 kHz). Arrow head, residue of artifact after the subtraction from neural recordings; double arrow, nerve activity evoked from A fiber

**Figure 2:**
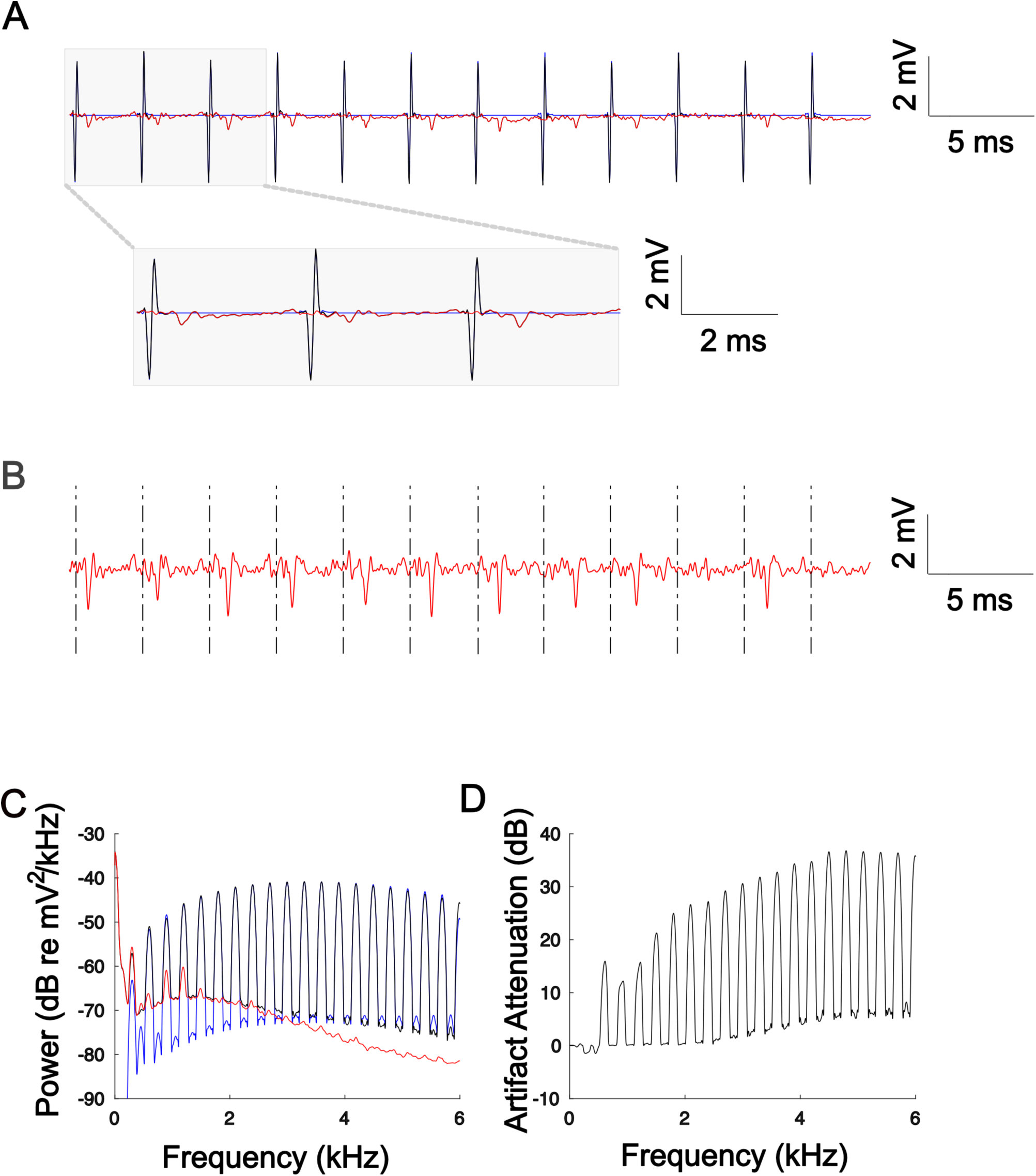
Artifact removal during bilateral cochlear implant stimulation and concurrent extracellular recordings in rat inferior colliculus. (A) Raw neural recordings (black) and predicted artifacts (blue) are largely overlapped. The cleaned neural recording trace (red) show no visible signs of artifact signals. (B) Zoomed version of the cleaned neural recording signal. (C) Power spectrum of the neural recording before (black) and after (red) artifact removal. The predicted artifact spectrum (obtained as the cross spectrum between recording trials) is shown in blue and largely overlaps the recorded spectrum prior to artifact removal (black). (D) Artifact attenuation ratio.

**Figure 3:**
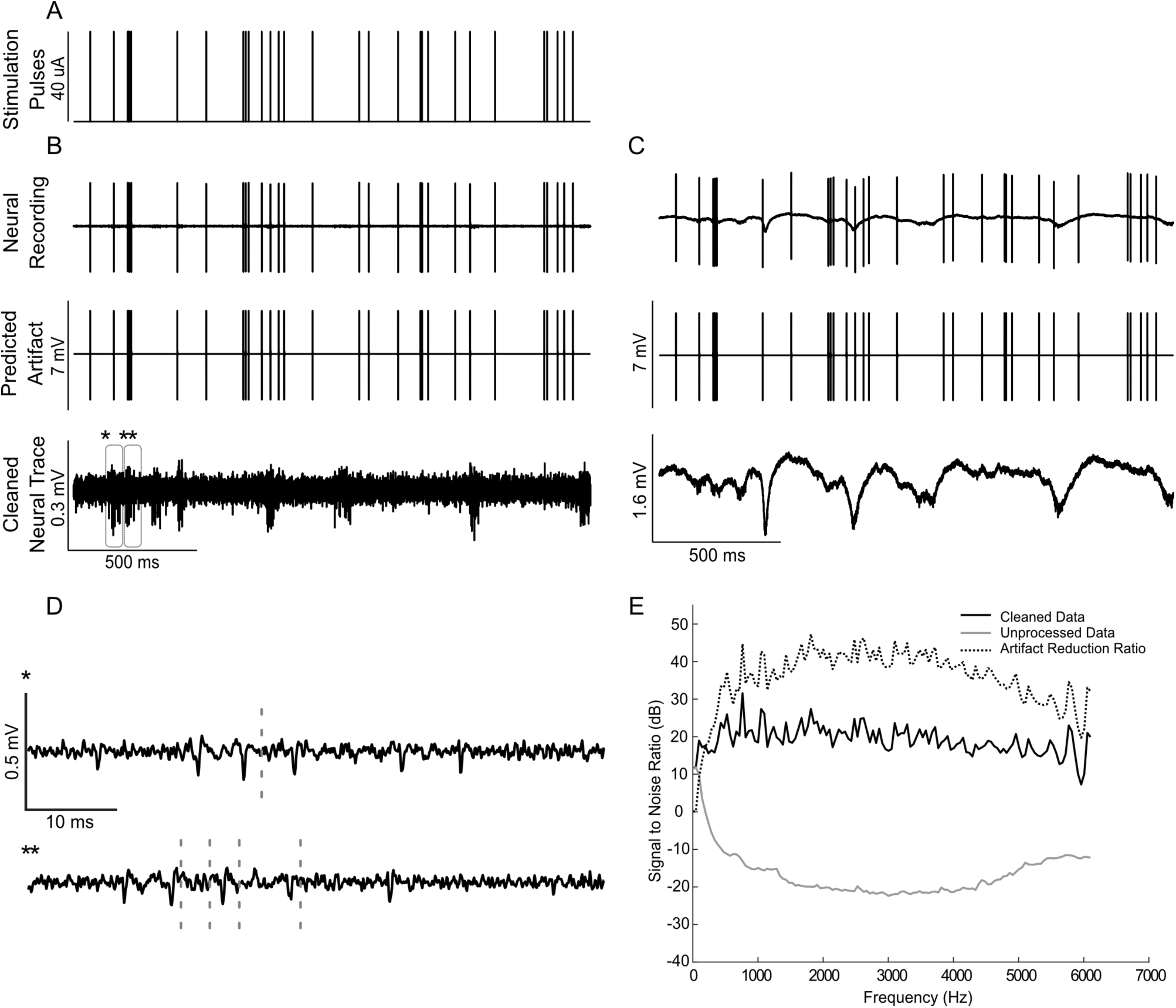
Artifact removal during a single channel electrical stimulation of the auditory midbrain and concurrent recording in auditory cortex. (A) Random Poisson distributed pulse sequence (average pulse rate of 16 Hz) were delivered to an electrode in auditory midbrain of a rat. Highpass filtered (B) and raw (C) neural recordings from a cortical electrode are dominated by the electrical artifacts (top). The estimated Wiener filters are used to predict the recorded artifacts (middle panels). Subtracting the artifacts from the neural recordings yields noise reduced estimates of the neural activity (bottom). (D) Zoomed sample waveforms showing the filtered extracellular signals after artifact subtraction (* and ** from panel B, bottom). (E) Signal to noise ratio prior to and after subtraction of the predicted artifacts (gray and black curve, respectively). The artifact reduction ratio is superimposed on the same panel (dotted curve).

**Figure 4:**
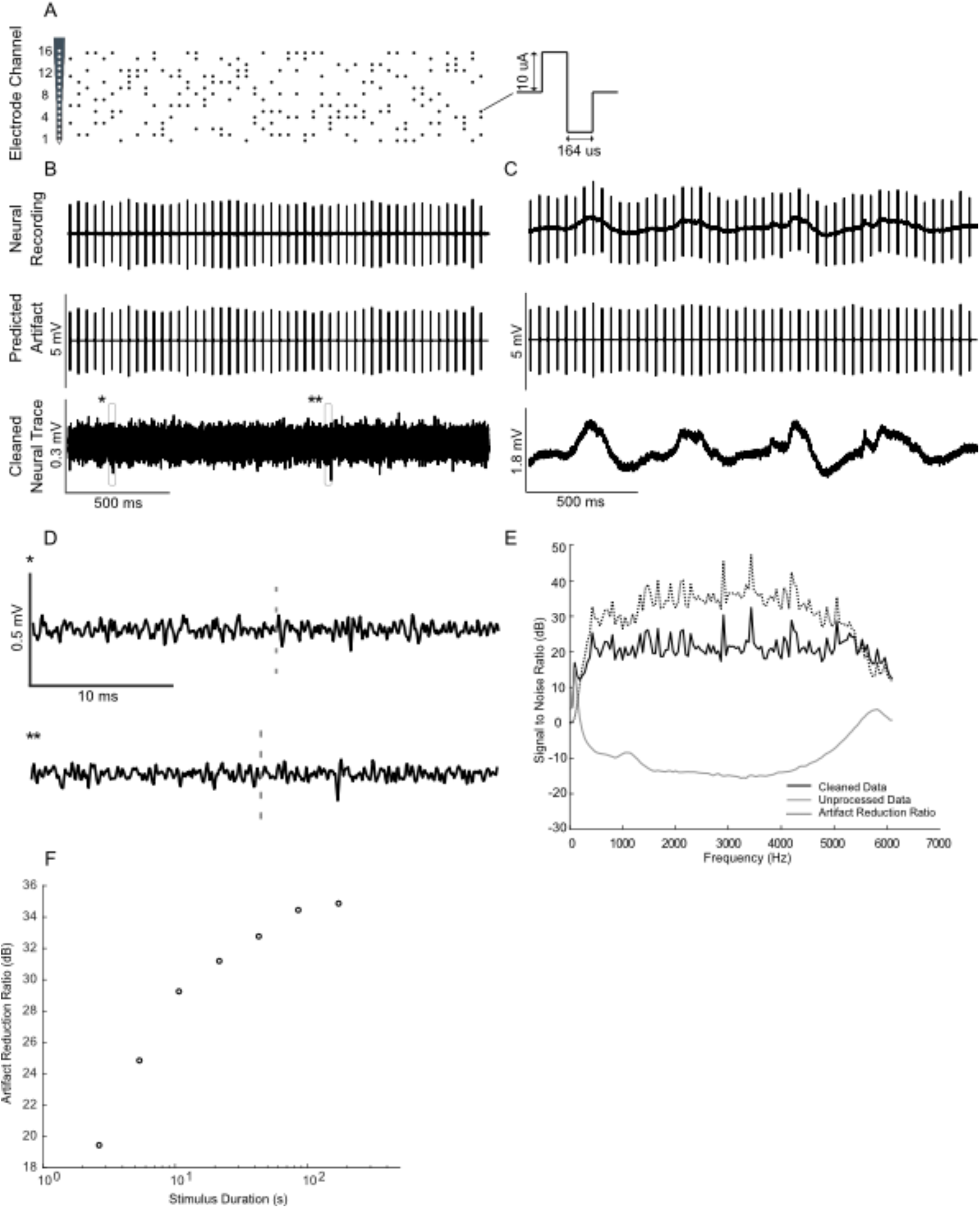
Artifact removal during high throughput multi-site electrical stimulation. (A) Spatiotemporal pulse sequence applied to a 16-channel probe placed in the auditory midbrain of a rat. Highpass filtered (B) and raw (C) neural recordings from a cortical electrode are dominated by the electrical artifacts (top). The estimated multi-channel Wiener filters are used to predict the recorded artifacts (middle panels). Subtracting the artifacts from the neural recordings yields noise reduced estimates of the neural activity (bottom). (D) Zoomed sample waveforms showing the filtered extracellular signals after artifact subtraction (* and ** from panel B, bottom). (E) Signal to noise ratio prior to and after subtraction of the predicted artifact is superimposed on the same panel (gray and black curve, respectively). The artifact reduction ratio is superimposed on the same panel (dotted curve). (F) Enhancement in artifact reduction ratio by increasing the electrical stimulus duration. The stimulus length is varied between 2.7-171.8 s in octave steps.

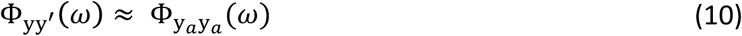

so that CSD between trials approximates the artifact noise spectrum. Furthermore, we note that for sufficiently long recordings 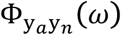 and 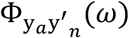 yield identical spectrum estimates on average and that 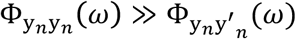 as a result of neural trial variability between trials. Thus, the neural signal spectrum can be approximated by subtracting the PSD from the CSD

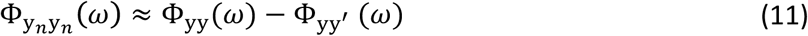

The signal to noise ratio is then approximated by

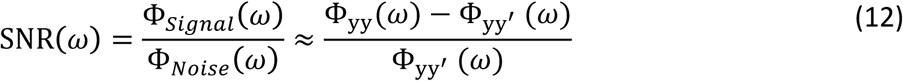

In the above, all cross and power spectral density estimates were obtained using a Welch average periodogram and a Kaiser window (*β* = 5, *N* = 256 time samples or 21 ms). To confirm the validity of the approximations used to derive Eqn. 12, we also estimated the SNR using an artifact free neural recording segment. Φ_*signal*_ (*ω*) was estimated by collecting 15 second neural trace without any electrical stimulation, which we then used to estimate the signal spectrum. We also estimated the noise spectrum directly from the Wiener filter predicted artifacts. Both procedures produce quantitatively similar results when compared to the original estimates (within ∼3 dB) confirming the approximations used to derive Eqn. 12.

### Artifact Reduction Ratio

In addition to defining the SNR metric, we also defined and measured an artifact reduction ratio (ARR). This metric quantifies the reduction in artifact power following artifact removal. It is defined as

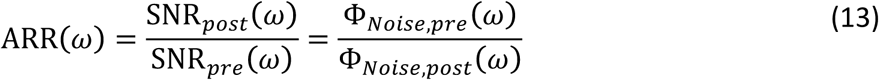

where SNR_*pre*_ (*ω*) is the SNR prior to artifact removal and SNR_*post*_ (*ω*) is the measured SNR after applying the artifact removal algorithm. Since the neural signal spectrum is unchanged by the artifact removal procedure, the above can also be estimated directly by taking the ratio of the noise spectrum prior to (Φ_*Noise,pre*_ (*ω*)) and post removal of the artifact (Φ_*Noise,post*_ (*ω*)). We note that the ARR metric is well defined for all frequencies whenever the electrical stimulation was delivered aperiodically since, in that case, the signal and noise spectrum take on continuous values at all frequencies (e.g., electrical stimulation of IC examples). However, for periodic electrical stimulation such as for the cochlear implant study (e.g., electrical stimulation periodically at 300 Hz), the electrical stimulation produced periodic artifacts with harmonic components in the signal spectrum at multiples of the stimulation frequency. Thus, the signal spectrum and hence the ARR contains signal components only at harmonics of the stimulation frequency and are thus well defined only at these components.

### Mouse Sciatic Nerve Recordings

#### Extracellular Recordings from Mouse Sciatic Nerve

All procedures were approved by the University of Connecticut Institutional Animal Care and Use Committee. Sciatic nerves of male C57BL/6 mice (6–8 weeks, Taconic, Germantown, NJ) were harvested for simultaneous extracellular recordings from teased nerve filaments as detailed previously [24]. Mice were anesthetized by isoflurane inhalation, euthanized by exsanguination from perforating the right atrium, and perfused through the left ventricle with oxygenated Krebs solution (in mM: 117.9 NaCl, 4.7 KCl, 25 NaHCO_3_, 1.3 NaH_2_PO_4_, 1.2 MgSO_4_, 2.5 CaCl_2_, and 11.1 D-glucose). Bilateral sciatic nerves of ∼30 mm long were harvested from their proximal projection to the L4 spinal cord to their distal branches innervating gastrocnemius muscles and transferred to a custom-built chamber perfused with oxygenated Krebs solution at 30 °C. The distal end of the sciatic nerve (∼5 mm) was gently pulled into a recording compartment filled with mineral oil and carefully split (i.e., teased) into fine neural filaments (∼25 μm thick) for extracellular recordings of action potentials. Extracellular recordings from multiple teased nerve filaments were conducted by a custom-built 5-channel electrode array consisting of micro-wires deployed parallel to each other with ∼150 μm clearance as described previously [24]. Action potentials were evoked at the un-teased end of the sciatic nerve using a platinum-iridium electrode (FHC Inc., ME). Multichannel recordings were digitized at 25 kHz, band-pass filtered (300 – 3000 Hz) and stored on a PC using an integrated neural recording and stimulating system (IZ2H stimulator, PZ5-32 neurodigitizer and RZ5D processor, TDT, Alchua, Florida, US).

#### Stimulation and Artifact Removal During Sciatic Nerve Recordings

We used the Wiener-Hopf equations defined above (Eqn. 3) to derive the impulse response of the artifact prediction filters during a sciatic nerve stimulation protocol. Electrical current inputs were delivered using a sub-and supra-threshold stimulation protocol consisting of a 120-s long low-frequency stimulations (0.5Hz, 0.2ms duration, cathodal current) with six ascending amplitudes (10, 20, 40, 80, 160, 320 μA; 10 stimuli per amplitude condition). Only recordings for which the stimulation current levels were sub-threshold (did not produce nerve activity) were used to derive the artifact prediction filters. These filters were then used to predict the stimulation artifacts for supra-threshold recordings. The predicted artifacts were then subtracted from the neurophysiological recordings to isolate the supra-threshold nerve response.

#### Validation of the Artifact Removal via Chemical Nerve Block

To quantify the efficiency of artifact removal via the Wiener filter artifact removal method, we used a non-selective sodium channel blocker (lidocaine) for several reasons. First, lidocaine treatment prevents action potential generation. The lidocaine treatment procedure thus allows us to isolate the artifact signal in the absence of background neural activity. This is useful for validating the accuracy of the artifact prediction since there is no confounding neural activity. Second, the prediction filters obtained during lidocaine treatment were also used to predict and remove the artifacts obtained in the absence of lidocaine treatment during supra-threshold stimulation. Thus, the approach allows us to isolate supra-threshold nerve activity and serves as cross validation to establish the reproducibility of the procedure across conditions, which is expected theoretically (with and without lidocaine; sub and supra threshold).

A bronze tube (4×4 mm cross section) was placed over the sciatic nerve to isolate a small segment of the nerve trunk (∼4mm) for lidocaine application. On both edges are small notch holes to allow nerve trunk to go through, which were lined with petrolatum to prevent solution exchange between inside and outside the bronze tube. Krebs solution inside the bronze tube was replaced with lidocaine (2% concentration, ∼0.2 ml) for 5 mins, and then the bronze tube was removed for bath washout for 15 min. Supra-threshold stimulation was conducted before, immediately after 5 mins lidocaine application and further after 15 mins of wash period.

The efficacy of the Wiener filter artifact removal procedure was estimated with the ARR as defined above (Eqn. 13). The ARR was estimated using the lidocaine treated data since the noise spectrum could be estimated directly from this data prior to and subsequent to artifact removal (i.e., there is no neural signal). Figure 1 displays the average ARR of 40 axons recorded across 300 −3000 Hz frequency range with an average artifact reduction ratio of 22.8 dB.

### Bilateral Cochlear Implant Stimulation in Rats

To illustrate the artifact removal during CI stimulation, we use example data from Wistar rat which was neonatally deafened by daily intraperitoneal (i.p.) injections of 400 mg/kg kanamycin from postnatal day 9 to 20 [25,26]. The animal was part of a study designed to determine factors governing sensitivity to binaural cues delivered via direct, intracochlear stimulation similar to that used in clinical CI devices. These data were obtained at the City University of Hong Kong, using procedures licensed by the Department of Health of Hong Kong (license number 16-52 DH/HA&P/8/2/5) and approved by the local ethical review committee. All surgical procedures, including CI implantation and craniotomy, were performed under anesthesia, which was induced with an i.p. injection of ketamine (80 mg/kg) and xylazine (12 mg/kg) and maintained by continuous i.p. infusion of ketamine (17.8 mg/kg/h) and xylazine (2.7 mg/kg/h) in 0.9% saline solution at a rate of 3.1 ml/h, and the animal’s body temperature was maintained at 38°C using a feedback-controlled heating pad (RWD Life Sciences, Shenzhen, China). The cochlear implantation methods are described in detail in [25,27]. In short, two to four rings of an eight channel intracochlear electrode carrier (ST08.45, Peira, Beerse, Belgium) were inserted through a cochleostomy in the medio-dorsal direction into the middle turn of both cochleae. The tip electrode ring of each intracochlear array was used to deliver electrical stimuli, while the second, adjacent electrode served as ground. Electrical stimuli were generated using a Tucker Davis Technology (TDT, Alachua, Florida, US) IZ2MH programmable constant current stimulator (TDT, Alachua, Florida, US) running at a sample rate of 24414 Hz. To verify that the cochlear implantation was successful and yielded symmetric evoked responses at comparatively low thresholds (typically less than 100 µA peak) in each ear, electrically evoked auditory brain stem response thresholds were measured for each ear individually. This was done by recording scalp potentials with subcutaneous needle electrodes implanted over the vertex and each bulla, averaged over the presentation of 400 individual biphasic electrical stimulus pulses.

A craniotomy was then performed bilaterally of the central cranial suture, just anterior to lambda, and a single-shaft, 32-channel silicon array electrode (ATLAS Neuroengineering, E32-50-S1-L6) was inserted stereotaxically into the IC through the overlying occipital cortex using a micromanipulator (RWD Life Sciences). Extracellular signals were recorded at a rate of 24414 Hz with a TDT RZ2 with a NeuroDigitizer headstage and BrainWare software. Neural tuning to interaural time differences (ITDs) of binaurally delivered pulse trains was then measured by recording extracellular responses of IC neurons to 200 ms trains of biphasic electrical pulses (duty cycle: 40.96 µs positive, 40.96 µs at zero, 40.96 µs negative), with peak pulse amplitudes approximately 6 dB above neural response thresholds and a rate of 300 pulses per second.. The recordings typically exhibited short response latencies (≈ 3-5 ms), which indicates that they probably come predominantly from the central region of IC.

### Electrical Stimulation of Auditory Midbrain and Cortical Recordings in Rat Auditory Cortex Surgical Procedures

All procedures were approved by the Institutional Animal Care and Use Committee of the University of Connecticut. Recordings were obtained from right cerebral hemisphere of adult male Brown Norway rats. Anesthesia was induced with ketamine and xylazine and maintained throughout the surgery and recording procedures. Depth of anesthesia was monitored using pedal reflex, heart rate, and blood oxygen saturation (SpO2) measured by a pulse oximeter. A heating pad was also used to maintain the animal’s body temperature at 37.0 ± 1.0 °C. Craniotomies were performed over the temporal cortex to make both cortex and inferior colliculus regions accessible. Dexamethasone and atropine sulfate were administered to reduce cerebral edema and secretions in the airway.

#### Electrophysiology

16-channel acute neural recording probes (NeuroNexus 5 mm probe; 16-linear spaced sites with 150 um separation; site impedance ∼100 *K*Ω) were used to record neural activity and also to deliver electrical stimulation to the inferior colliculus (IC). Stimulating and recording probes were grounded to the animal’s neck muscle and the eye bars holding the animal in place, respectively [28]. The probes were inserted with a high precision LS6000 microdrive (Burleigh EXFO). A 4-channel acute single-shank recording probe (Qtrode, NeuroNexus Inc; 5 mm shank length, tetrode with 25 um site separation; site impedance ∼1-3 *M*Ω) was simultaneously inserted into auditory cortex (AC). Penetration sites were chosen within the depth range of cortical layer IV where AC receives its inputs from auditory thalamus. A sequence of pure tones with varying frequency and attenuation was initially played to the animal’s left ear (contralateral to the brain opening) and brain responses were recorded to generate frequency response areas (FRA) to verify probes placements in the central nucleus of IC (CNIC) and AC.

Neural activity was recorded digitally at a sampling rate of 12 kHz using a PZ2 preamplifier and RZ2 real time processor (TDT, Alchua, Florida, US). Electrical stimuli were delivered to the IC electrode via the IZ2 stimulation module (TDT, Alchua, Florida, US). Electrical pulse sequences with amplitudes of either 40 µA or 10 µA were transmitted to a single electrode or independently across multiple electrode channels, respectively (see below for details). Neural activity was then recorded from the auditory cortical probe for the duration of each stimulus.

#### Single-Channel and Multi-Channel Electrical Stimulation

We first delivered Poisson-distributed biphasic pulse sequence during single channel electrical stimulation. A random sparse sequence of impulses with arrival time following Poisson point process and impulse rate of 16 Hz was first generated. The impulse sequence was convolved with a biphasic pulse (164 µs duration and 40 µA current amplitude) to produce the current waveform used for electrical stimulation.

For multi-site electrical stimulation, we delivered a random quad-pulse train sequence (RQP). The RQP sequence is generated by delivering biphasic pulses (164 µs duration and 10 µA amplitude) concurrently across 4 randomly chosen electrode channels every 40 ms yielding an average pulse rate of 100 pulses/s as illustrated in Fig. 4. This multi-site sequence produces a random spatio-temporal patterned set of pulses that are delivered across the 16-channel electrode array.

## RESULTS

We demonstrate the Wiener filter effectiveness at predicting and removing neural recordings artifacts during single and multi-channel electrical stimulation for both high-frequency spiking activity and low-frequency local field potentials (LFP) in a variety of recording modalities. The success of the artifact removal method is evaluated by comparing the residual artifacts across repeated stimulation trials and estimating neural recording SNR before and after removing artifacts.

### Single-Channel Electrical Stimulation of Sciatic Nerve

Monopolar stimulus pulses (0.2 ms duration, cathodal current, 0.5 Hz) with 6 current amplitudes (starting with subthreshold current amplitude, logarithmic scale of log2) were delivered to one end of sciatic nerve by a platinum iridium electrode. Evoked action potentials along with stimulus artifacts were recorded from 40 teased sciatic nerve filaments. Transfer functions for each stimulation-recording electrode pair were identified by Wiener-Hopf equation using the 120 s long subthreshold stimulus protocol via application of Lidocaine treatment (see Methods). This allowed us to measure isolated artifacts in the absence of neural activity and allowed us to derive artifact prediction filter estimates. In subsequent supra-threshold stimulation and recordings experiments, the derived filters were used to predict the artifact waveforms by convolving with the suprathreshold electrical stimulation sequences. This procedure serves as form of cross-validation and further serves as an assessment of linearity. The predicted waveforms as illustrated in Fig. 1 (B and E in black) were then subtracted from the neurophysiological recordings. As can be seen in Fig. 1 (C and F in purple) the artifact amplitudes were substantially reduced upon subtraction. As a control, we also obtained recordings following the application of lidocaine which blocks action potentials generation so that the recorded signals consisted of pure stimulus artifacts as shown in Fig. 1 (B and E in red). This post-lidocaine artifact signal was then subtracted from the original recordings (pre-lidocaine) which allows us to isolate the neural response component (Fig. 1 C and F in green). As exemplified in Fig. 1 (C and F), the lidocaine-subtracted neuronal components is almost identical to the isolated neural signals obtained by subtracting the Wiener filter predicted artifacts. As shown in Fig. 1 G, the average artifact reduction ratio by implementing the Wiener filter predication is on average 22.8 dB across the frequency range (300 – 3000 Hz).

### Bilateral Cochlear Implant Stimulation

The artifact removal procedure was also tested with high-rate bilateral cochlear implant stimulation in rat while concurrently recording from a silicon array electrode implanted in the IC. Biphasic electrical pulse sequences were delivered at a pulse rate of 300 Hz synchronously to both ears, at different interaural delays (see Methods). An example raw recorded waveform from one IC electrode channel is shown in Fig. 2A (black), along with the predicted artifact waveform (blue). As can be seen, the predicted artifacts signals are largely superimposed and are visually indistinguishable from the artifacts on the neural recordings. Synchronized action potentials are observed immediately following the delivery of electrical stimulus current pulses. Upon subtracting the predicted artifact (blue) from the neural trace (black), the cleaned neural trace is exceptionally clean with no evident sign of stimulation artifacts and no evident sign of waveform distortions (Fig. 2A & B, red). Spectral analysis of the recorded signal prior to (Fig. 2C, black) and after artifact removal (Fig. 2C, red) confirms a substantial reduction in the artifact size. The original artifact spectrum has harmonic component with 300 Hz fundamental (blue) which dominates the original recording (black). Upon removal of the predicted artifact, there is a substantial reduction in the artifact components (red). Overall, the average artifact attenuation at harmonics of the stimulation frequency is 27 dB (between 300-5000 Hz; Fig. 2D).

### Single-and Multi-Channel Electrical Stimulation in Auditory Midbrain

We also tested the artifact removal procedure by delivering random biphasic electrical pulse sequences (Poisson distributed pulse intervals, 164 µs pulse duration, and 40 µA current amplitude, Fig. 3A) to an auditory midbrain electrode while neural activity was concurrently recorded from rat auditory cortex. As can be seen in Fig. 3B and C, the extracellular neural activity (Fig. 3B, highpass filtered above 300 Hz) and the corresponding unfiltered recordings (Fig. 3C, unfiltered) both contain stimulation artifacts that are substantially larger than the target neural signals.

We numerically estimated a digital Wiener filter (N=40 order) to predict and subsequently remove the electrical stimulation artifacts (see Methods). Fig. 3B and C show the raw cortical recordings (top panels), the predicted artifacts (middle panel) and cleaned neural traces obtained by subtracting the predicted artifacts from the raw recordings. The artifact prediction algorithm accurately predicts the timing and amplitude waveform of the electrical artifacts and, upon subtraction, the procedure successfully isolates either the extracellular waveforms or low-frequency local field potentials in the neural signal. Magnified traces of the extracellular recordings (marked by * and **) are presented in Fig. 3D to show the cleaned neural recordings at a higher resolution. Notably, the algorithm is able to subtract the artifacts that occur in the vicinity of neural spiking with no visible signs of neural waveform distortions.

Performance metrics of the artifact prediction and subtraction algorithm for this recording is shown in Fig. 3E (applied to the broadband unfiltered signal). The signal-to-noise ratio of the original recorded waveform varies with frequency but is generally in the order of −10 to −20 dB. Upon subtracting the predicted artifact, the cleaned SNR is ∼20 dB with an average SNR enhancement ranging between 30 to 45 dB (average = 39 dB between 300-5000 Hz). Thus, there is a marked reduction in the artifact size and, as seen in the zoomed neural recordings, there are no visible distortions created by the subtraction algorithm.

We also successfully used the artifact removal during high throughput multi-channel electrical stimulation (16 stimulation channels) of the auditory midbrain. Random pulse sequences (100 pulses/s) were delivered to the 16-channel auditory midbrain array (Fig. 4A; 10 µA pulses delivered across four randomly chosen electrode channels at a time) while recording from an auditory cortex electrode. For this multi-stimulation site configuration, we numerically estimated the digital filters that predict the artifacts generated by each of the electrical stimulation channel. Filtered and unfiltered neural recordings, predicted artifacts, and the cleaned neural traces are depicted for both the filtered (Fig. 4B) and unfiltered (Fig. 4C) data. As for the single channel electrical stimulation, the artifact prediction filter is able to accurately predict the measured artifacts during multi-channel electrical stimulation, resulting in minimal distortion of the extracellular signals or the local field potentials. Prior to removing the artifact, the SNR for this recording dips to approximately −15 dB at ∼3kHz (Fig. 4E). Following artifact removal the SNR hovers around ∼20 dB with an overall improvement in the range of 30-45 dB across the frequency range (average artifact reduction ratio=33.5 dB from 300-5000 Hz; Fig. 4E).

### The Impact of Data Length on Artifact Removal Quality

As seen from different examples, there are some discrepancies in the artifact reduction ratio between the different recordings. For instance, the sciatic nerve recording ARR was ∼ 23 dB whereas for the cochlear implant and cortical recordings of Fig. 2-4 the ARR was somewhat higher (∼30 to 40 dB). This discrepancy is in part accounted by the quality of the estimated artifact prediction filters, which is expected to depend on the length of the recording experiment and the number of pulses delivered. For instance, the sciatic nerve experiments used slow rate pulse sequences (0.5 pulses/s) of relatively short duration and thus relatively few artifact measurements (tens of pulses total) were used to estimate the artifact prediction filters which likely resulted in the low ARR. This contrast the auditory midbrain and cortical recordings of Fig. 2-4, where longer sequences were used and pulses were delivered at a much higher rate (Fig. 2, 300 pulses/s; Fig. 3, 16 pulses/s; Fig. 4, 100 pulses/s) which resulted in a much higher number of artifact measurements for the filter estimation. The impact of the estimation data length (or equivalently number of artifact pulses used to estimate the filters) on the quality of the algorithm are shown for the auditory cortex recording of Fig. 4 (F). The recorded data was portioned into segments of a fixed duration (2.7 – 172 s; corresponding to ∼270–17,200 artifacts) and the filters were re-estimated using the partitioned data followed by the artifact prediction and removal procedure. As expected, the ARR improves with increasing estimation data length, or equivalently the number of artifacts used to estimate the filters, with an average improvement of ∼2.5 dB per doubling of the data length.

## DISCUSSION

We have developed a linear predictive filtering methodology that can be used to predict and subtract electrical stimulation artifacts from neural recording data in a wide range of applications and recording modalities, including high rate and multi-channel electrical stimulation. The method was validated in three different neural stimulation settings: single-channel stimulation of sciatic nerve axons, bilateral (two-channel) CI stimulation and multi-channel stimulation of the auditory midbrain. In these experiments the procedure reduces the artifacts size by 20-40 dB without visibly distorting the surrounding neural activity, although further improvements can be gained as additional data is acquired. Furthermore, as demonstrated, the procedure is efficient requiring only modest sized datasets (10-100 s) to estimate the artifact prediction filters and to obtain noticeable improvements in signal quality.

In contrast to conventional artifact removal procedures, which are designed to be blind to the artifact generation source, our method capitalizes on the fact that artifact source currents are actually known when stimulating neural tissue. Furthermore, the procedure capitalizes on linearity, since electrical artifacts arise through passive linear tissue conduction and capacitive coupling at the electrode interface [19]. As such, the Wiener filter approach can accurately model the linear transformations between the stimulation sources and recording electrodes, which allows us to accurately predict and subsequently subtract recording artifacts from the neural recordings. Conventional procedures based on template subtraction are often not able to eliminate artifacts since finding a template that matches the shape of all artifact waveforms is not always possible [5,6,8]. Such is the case for multi-site stimulation or whenever the electrical stimulation currents have varied amplitudes or shapes. Furthermore, the extracted templates can contain neural fluctuations which causes distortion in neural traces [6]. Other established artifact removal procedures, such as those utilizing independent component analysis [11-16], are designed on an assumption of independence between neural and artifact sources, which is not entirely satisfied since neural activity can synchronizes to electrical stimulation. Thus, some of the estimated artifact components can contain both neural activity and artifacts, which can distort and eliminate relevant neural signals. Finally, while recent developments have had success in removing artifacts from multiple sources for spike sorting neural data [17], these model-based approaches place constraints on the statistics of the data and do not make use of the known stimulation currents which, as shown here, provides substantial predictive power of the generated artifacts.

As seen for the different neural stimulation and recording modalities tested, the quality of artifact reduction varied by ∼ 20 dB (ARR range =∼20-40 dB). Although the mouse sciatic nerve example yielded the lowest improvement with only a modest reduction of the artifact size by ∼20 dB, we point out that the artifact prediction filters for that set of neural recordings were generated with short recording segments containing only tens of recorded artifacts. This contrast the neural recordings in IC and AC, which used high-rate stimulation and longer recordings, thus allowing us to train the artifact prediction filters with substantially more data and many more artifact measurements. As shown in Fig.4, the quality of the artifact removal can be improved substantially if a longer data and more artifacts are used for the filter estimation yielding approximately a 2.5 dB improvement for each doubling of the data length for that example (or equivalently, doubling the number of induced artifacts). In practice, Wiener filters can be estimated and implemented iteratively using solutions that update the coefficients as additional data is acquired [20] and such an approach will be investigated in a future study. In theory, it allows the filter quality to improve as additional data is acquired and would also allow estimation of the filter coefficients in real-time. Such iterative implementations would also be useful to localize the estimated filters in time, which could be valuable for long term chronic recordings that may exhibit nonstationary behavior (e.g., due to changing electrode impedance over days or movement of electrodes etc.).

The artifact removal algorithm was able to remove stimulation artifacts without noticeable distortion of the neural spiking activity, as demonstrated in the recordings from mouse sciatic nerve axons. Lidocaine, as a non-selective sodium channel blocker, was used to block neural activity leaving only artifact signals, which allows us to validate the artifact prediction algorithm. As shown in Fig. 1 panel E, the predicted artifact closely matched the artifact after blocking neural activity with lidocaine. This was true even though the artifact prediction filters were generated using sub-threshold current stimulation and subsequently tested using much higher supra-threshold currents. Fast-conducting A-fiber in Fig. 1 panel F was isolated after subtracting the predicted artifact without any distortion, which was verified by the same action potential preserved after subtracting lidocaine treated artifact. This allowed us to isolate neural activity from fast-conducting A-fibers which are typically embedded within the artifact signals. Together, the combined use of lidocaine treatment and the use of sub-and supra-threshold current stimulation to separately estimate and test the algorithm provides validation to the assumption that artifact generation is largely a linear process between stimulation and recording sites.

The technique can also be applied in a variety of electrical stimulation settings, including both low-frequency (pulse rate of 0.5 Hz in sciatic nerve stimulation) and high-frequency (pulse rate of ∼300 Hz in CI) electrical stimulation, and is capable of isolating the desired neural signals for both single-unit (single-channel stimulation of sciatic nerve axons/inferior colliculus) and multi-unit (Bilateral CI stimulation and 16-channel stimulation of inferior colliculus) electrical stimulation modalities. Furthermore, the algorithm was able to successfully isolate both high-frequency spiking activity and continuous LFPs. Overall, this methodology has potential for broad range of applications requiring concurrent neural stimulation and neural recording from multiple channels. The Wiener filter estimation and prediction approach is well established and computationally efficient requiring relatively short recordings to estimate the artifact prediction filters, low computational resources, and does not require specialized hardware. Hence, the approach can be easily adapted for real-time applications and applications requiring real-time assessment of neural function and behavior.

## Acknowledgements

The authors would like to thank Dr. A. Buck and Ms. Kongyan Li for assistance with the acquisition of the binaural cochlear implant data presented in this manuscript.

